# Impaired HSF1 transactivation drives proteostasis collapse and senescent phenotype of IPF lung fibroblast

**DOI:** 10.1101/2020.04.10.036327

**Authors:** Karina Cuevas-Mora, Willy Roque, Dominic Sales, Jeffrey D. Ritzenthaler, Edilson Torres-Gonzales, Andrew J Halayko, Ivan O. Rosas, Jesse Roman, Freddy Romero

**Affiliations:** Department of Medicine, Division of Pulmonary, Allergy and Critical Care and the Center for Translational Medicine; the Jane & Leonard Korman Respiratory Institute, Philadelphia, PA, US; Department of Medicine, Rutgers University, New Jersey Medical School, Newark, NJ. US; Department of Physiology and Pathophysiology, University of Manitoba, Winnipeg, MB Canada R3E3P4; Pulmonary, Critical Care and Sleep Medicine, Baylor College of Medicine, Houston, TX. US

**Author notes:** To whom correspondence should be addressed: Department of Medicine, Division of Pulmonary, Allergy and Critical Care and the Center for Translational Medicine and Jane & Leonard Korman Respiratory Institute, Thomas Jefferson University, 1020 Locust Street, JAH 354, Philadelphia, PA 19107, USA. These authors contributed equally to this work (Karina Cuevas-Mora, Willy Roque, Freddy Romero). Author Contributions: Data analysis and interpretation, K.C.M., W.R, D.S., A.J.H., I.O.R, J.R., F.R. Writing. K.C.M, W.R., J.D.R., I.O.R., J.R. and F.R. Execution of experiments, D.S., W.R., K.C.M., E.T.G., J.D.R., F.R. Planning. K.C.M., W.R, D.S., F.R. Funding, F.R. Acquisition of data: D.S. W.R., K.C.M., F.R. All authors participated in the intellectual revision and approved the final version of the manuscript.

**Keywords:** Heat shock proteins, proteostasis, cellular senescence, idiopathic pulmonary fibrosis, chronic obstructive pulmonary disease, sumoylation

## Abstract

Loss of proteostasis and cellular senescence are key hallmarks of aging. Recent studies suggest that lung fibroblasts from idiopathic pulmonary fibrosis (IPF) show features of cellular senescence, decline in heat shock proteins (HSPs) expression and impaired protein homeostasis (proteostasis). However, direct cause-effect relationships are still mostly unknown. In this study, we sought to investigate whether the heat shock factor 1 (HSF1), a major transcription factor that regulates the cellular HSPs network and cytoplasmic proteostasis, contributes to cellular senescence in lung fibroblasts. We found that IPF lung fibroblasts showed an upregulation in the expression of various cellular senescence markers, including β-galactosidase activity (SA-β-gal) staining, the DNA damage marker γH2Ax, the cell cycle inhibitor protein p21, and multiple senescence-associated secretory proteins (SASP), as well as upregulation of collagen 1a1, fibronectin and alpha-smooth muscle actin (α-SMA) gene expression compared with age-matched controls. These changes were associated with impaired proteostasis, as judged by an increase in levels of p-HSF1^ser307^ and HSF1^K298 sumo^, downregulation of HSPs expression, and increased cellular protein aggregation. Similarly, lung fibroblasts isolated from a mouse model of bleomycin-induced lung fibrosis and mouse lung fibroblast chronically treated with H_2_O_2_ showed downregulation in HSPs and increased in cellular senescence and SASP markers. Moreover, sustained pharmacologic activation of HSF1 increased the expression of HSPs, reduced cellular senescence markers and effectively reduced the expression of pro-fibrotic genes in IPF fibroblast. Our data provide evidence that the HSF1-mediated proteostasis is important for driving lung fibroblasts toward cellular senescence and a myofibroblast phenotype. We postulate that enhancing HSF1 activity could be effective in the treatment of lung fibrosis.

## INTRODUCTION

Idiopathic pulmonary fibrosis (IPF) is a progressive lung disease that impairs quality of life and proves fatal in the majority of sufferers within 3-4 years of diagnosis. It is estimated that at the time of disease presentation, two-thirds of the patients with IPF are over the age of 60 years [1-3]. Currently, there are a limited number of marginally effective treatment options for patients with IPF and related progressive forms of fibrotic lung disease, emphasizing the need for further mechanistic insight and translational progress [4, 5].

Although the etiology of IPF is not fully understood, it is now generally accepted that fibroblast activation plays a critical role in the development and progression of IPF [6, 7]. While there is mounting evidence to support an important contribution of mitochondrial dysfunction and cellular senescence in IPF [8-10], recent studies point to a pathogenic role for impaired cellular proteostasis in promoting fibrotic responses. These studies suggest that impaired HSP70 expression induces fibrotic remodeling through a variety of mechanisms including the activation of an apoptotic resistant phenotype and inflammatory pathways [11-13].

Cellular senescence is a physiological state in which cells lose their capacity to proliferate, but maintain a high rate of metabolic activity and secrete a multitude of factors [14]. Cellular senescence has been mechanistically linked to several common pathological conditions, including chronic obstructive pulmonary disease and IPF [15-17]. Senescent lung fibroblasts exhibit myofibroblast-like characteristics, including increased levels of α-smooth muscle actin (α-SMA), collagen 1a1 and fibronectin. Moreover, recent studies have shown that age-dependent accumulation of senescent myofibroblast may block fibrosis resolution following bleomycin exposure, and that targeting senescent cells for destruction can be quite effective in ameliorating experimentally induced respiratory conditions [18, 19], highlighting a potential role for anti-senescent approaches in the treatment of chronic lung disease.

Although induction of cellular senescence has been attributed to a variety of factors, such as telomere attrition or mitochondrial dysfunction [20, 21], decline in the proteostasis network (PN) has recently emerged as an important causal factor [22-24]. The PN is a nexus of pathways that act in concert to maintain the integrity of the proteome. Under proteotoxic stress, cells are known to elicit a cellular stress response that include the activation of autophagy, unfolded protein response (UPR), the 26S proteasome system and HSF1-mediated chaperones expression [25, 26]. Molecular chaperones function to promote efficient folding and target misfolded proteins for refolding or degradation. The expression and efficiency of the molecular chaperones declines with age, resulting in the accumulation of misfolded proteins, and in some cases in the development of age-related fibrosis [11, 12]. Thus, maintaining an active and efficient PN through the late stages of life could delay or prevent fibrotic diseases.

The HSF1 protein regulates the expression of HSPs to maintain cellular proteostasis [27, 28]. Under normal conditions, HSF1 exists in a predominantly monomeric and transcriptionally repressed state, which lacks the ability to bind the cis-acting heat shock elements located in the promoters of HSPs genes. HSP90 and HSP70 act as cellular repressor of HSF1 and play a major role in retaining HSF1 in a monomeric state. Induction of transcriptional activity by cellular stress then results in the conversion of HSF1 from an inactive monomer to a DNA-binding trimer [29, 30]. Despite its central role in stress resistance, disease and aging, the mechanisms controlling HSF1 transactivation remain incompletely understood. Initially, HSF1 was believed to cause cellular senescence only in situations in which cellular expression is diminished [31, 32]. In this scenario, reduced expression of HSF1 has been shown to cause senescence by eliciting a proteostasis collapse, thereby compromising the ability of cells to perform essential biological activities. However, it is now appreciated that increased post-transcriptional modification, induced via sumoylation, acetylation and phosphorylation can also contribute to impaired HSF1 activity. In these circumstances, it has been shown that modulation of HSF1 activity through Ser303/Ser307 phosphorylation and sumoylation at K298 are induced hyperactivity, thereby resulting in decline HSF1 transcriptional activation and increased oxidant-induced cellular damage [33, 34].

In this study, our objective was to determine whether HSF1 dysfunction contributes to cellular senescence in the IPF lung fibroblast. Using three different approaches: 1) an *ex vivo* model, in which fibroblasts were isolated from the lungs of IPF and controls donors; 2) an animal model, in which fibroblasts were isolated from the lungs of animals treated with bleomcyin biweekly; and 3) an *in vitro* model, in which mouse fibroblasts are exposed to a chronic low dose of H_2_O_2_. Here we assessed whether HSF1 expression and function were altered in senescent fibroblast cells. Then, after confirming that impaired transcriptional activation was present in all models, we tested whether strategies aimed at restoring HSF1 homeostasis in cultured fibroblasts could alleviate the senescent and myofibroblast phenotype.

## METHODS

### Animals

C57B/6J mice (8-10 weeks old) were purchased from the Jackson Laboratory (Bar Harbor, ME) and housed in a pathogen-free animal facility at Thomas Jefferson University. Throughout the study period, mice were maintained on a standard chow diet (13.5% calories from fat, 58% from carbohydrates, and 28.5% from protein) and permitted to feed *ad libitum*. Prior to the initiation of any study, the Institutional Animal Care and Use Committee at Thomas Jefferson University approved all animal protocols.

### Human subjects

Lung fibroblasts were harvested from the lungs of twelve subjects. Six IPF lung specimens were provided by Dr. David Nunley (previously in the Lung Transplantation Program, Department of Medicine, at the University of Louisville; these samples were obtained from explants of six IPF patients undergoing lung transplantation at the University of Louisville, Louisville, Kentucky. All patients provided written consent and the study was approved by the University of Louisville Hospital Ethics Committee and the Institutional Research Committee (no number included). Non-IPF lung fibroblasts were generously provided by Dr. Andrew Halayko (Department of Physiology & Pathophysiology, Pediatrics and Child Health, and Internal Medicine, University of Manitoba, Winnipeg, MB, Canada). They were isolated from the uninvolved periphery of the organ of six lung cancer patients. Fresh tissues were transported immediately to the laboratory for isolating fibroblasts according to procedures described before [35].

### Bleomycin-induced lung injury

Bleomycin sulphate (Enzo life science, BML-AP302-0050) was dissolved in phosphate buffered saline. Lung injury was induced by instilling 0.04U of bleomycin into the posterior oropharynx of anesthetized mice every other week for five doses [36]. The mice were observed following intubation to ensure complete recovery from anesthesia. Mice were euthanized 2 weeks after the last dose by exposure to carbon dioxide, and lungs were harvested for fibroblast isolation and histological preparations as detailed below and as previously described.

### Lung histology

The trachea was cannulated, and the lungs were inflated with 4% paraformaldehyde (Sigma, 1004969019) in PBS. Lungs were removed *en bloc* and immersed in the same fixative at 4°C for 18 h and then embedded in either paraffin or optimum cutting temperature (OCT), LAMB-OCT-USA. Hematoxylin and Masson’s trichrome stain (Millipore Sigma, HT15-1KT) were performed on four-µm-thick sections.

### Isolation of primary fibroblast cells

Primary lung fibroblasts were obtained from mouse lungs by cutting the lung parenchymal tissue into 1-mm sections. Tissue sections were washed twice in sterile PBS, resuspended in DMEM with 4.5 g/L glucose (ThermoFisher, 11995040), 10% FBS (ThermoFisher, 26140079), 1% antibiotic-antimycotic solution (100 U/mL penicillin G sodium, 100 U/mL streptomycin, 0.25 µg/mL amphotericin B), (Millipore Sigma, A5955) transferred to a tissue culture dish, and incubated in a humidified 5% CO_2_ incubator at 37°C for 1 to 2 weeks to allow fibroblasts to migrate out of tissue sections. Primary lung fibroblasts were between 2-5 passages when used in experiments [37].

### Cell culture and reagents

Mouse lung fibroblast were obtained from the American Type Culture Collection (ATCC, CCL-206). Cellular senescence was induced by exposing cells to 10 μM of H_2_O_2_ (Sigma-Aldrich, H3410) for five consecutive days and maintaining cells in culture for an additional 2 days. Cellular senescence markers were evaluated at day 7 after the initiation of H_2_O_2_ treatment.2-2(2-Benzoxazolyl)-3-(3-pyridinyl)-1-(3,4,5-trimethoxyphenyl)-2-Propen-1-one (A3), CAS496011-51.9, ChemBridge 7355596 was used at a concentration of 5 μM except where otherwise specified.

### Oxygen consumption measurements

Oxygen consumption rate (OCR) was measured using the Seahorse XFp Bioanalyzer (Seahorse Bioscience). In brief, lung fibroblasts were seeded at a concentration of 30,000 cells/well on XFp cell plates (Seahorse Bioscience, 103057-100) 24 hours prior to the initiation of studies. The concentration of FCCP (also known as trifluoromethoxy carbonylcyanide phenylhydrazone), antimycin A, and rotenone were 2μM, 0.75μM, and 1μM respectively. Pyruvate (10 mM) and glucose (25 mM) served as a substrate, in addition to ADP (4 mM). The final concentrations of injected compounds were as follows: port A, 2 μM rotenone; port B, 10 mM succinate; port C, 4 μM antimycin A; port D, 100 mM ascorbate plus 1 mM TMPD (tetramethyl-p-phenylenediamine). The protocol and algorithm for our XF-PMP assay were designed using wave 2.4 software.

### Senescence-associated beta-galactosidase (SA-β*-gal) detection*

SA-β-gal staining was performed using the β-galactosidase kit (Dojindo Molecular Technology, 1824699-57-1) according to manufacturer’s instructions. To prevent false positives, experiments were performed on cells at 70% confluence.

### Protein aggregation assay

A Proteostat aggresome detection kit (Enzo Life Sciences, ENZ-51035-0025) was used to characterize the aggregates and aggresomes in senescent lung fibroblast. All components of this kit were prepared according to the manufacturer’s instructions. Monolayer cells was grown on cell culture dishes. Cells were washed with 1X PBS and fixed with 4% paraformaldehyde for 10 min at room temperature. After washed with excess 1 X PBS, cells were permeabilized with 0.5% Triton X-100 (Millipore-Sigma, 9002-931), 3 mM EDTA (Sigma-Aldrich, 03690) in 1 X PBS for 30 min. Cells were again washed twice with 1X PBS and the ProteosStat dye was added at 1:2000 dilution for 30 min.

### Proteasome activity

Lung fibroblast were washed twice with cold PBS, collected with a rubber policeman and lysed in ice-cold proteasome buffer (10 mM Tris, pH 7.8, 1 mM EDTA, 5 mM MgCl_2_, 5 mM ATP). Cell debris was removed by centrifugation for 15 min at 14,000 x g and the supernatant was used for the assay. Chymotrypsin-like, trypsin-like and caspase-like proteasome activity was assayed at 37 °C using 10 µg of protein extracts in proteasome buffer in the presence of 100 μM Suc-LLVY-aminoluciferin (Succinyl-leucine-valine-tyrosine-aminoluciferin), Z-LRR-aminoluciferin (Z-leucine-arginine-arginine-aminoluciferin and Z-nLPnLD-aminoluciferin(Z-norleucine-proline-norleucine-aspartate-aminoluciferin), respectively (Promega Corporation, G8531). The reaction was monitored by fluorimetric measurements every 10 min for 1h (excitation 350 nm, emission 460 nm) in a Synergy HT Multi-Detection microplate reader. Proteasome activity was determined as the difference between the total activity of cell lysis and the remaining activity in the presence of 20 µM MG132 (Millipore-Sigma, M7449).

### RNA isolation and Real-time quantitative PCR (qRT-PCR)

Gene transcript levels were quantified by real-time PCR. Total RNA was isolated using RNeasy Mini-Kit (QIAGEN, 74104), according to the manufacturer’s instructions. A summary of primer sets manufactured by Integrated DNA Technologies can be found in supplemental Table 1. Relative gene expression levels were quantified with the Livak method (2-ΔΔCT) by normalizing expression of the gene of interest to the combined expression of two stable housekeeping genes, demonstrated to be unaffected by A3 (18S and Gadph), and calculating the relative fold differences between experimental and control samples. Assays were completed in technical triplicate.

### Western blot analysis

Fibroblast cells were lysed in ice cold buffer (PBS, 0.05% Tween 20, pH 7.4) containing protease inhibitors (Active motif, 37491) and phosphatase inhibitor (Active motif, 37493). After centrifugation (14,000 x g, 10 min, 4°C), the supernatant was collected. Fibroblast cell nuclear fraction was extracted using a commercially available kit (Active motif, 40010) according to the manufacturer’s instructions. Protein concentration was determined by Pierce™ BCA assay kit (ThermoFisher, 23225). Protein samples (20 μg) were solubilized in 4 × Laemmli sample buffer, heated at 95°C for 10 min, centrifuged at 3,000 g for 1 min, loaded on a 10% Tris-HCl-SDS-polyacrylamide gel and run for 1 h at 100 V. Protein was transferred to a nitrocellulose membrane (ThermoFisher, LC2000) and then blocked with Odyssey Blocking Buffer (Li-Cor Biosciences, 927-50000) for 1 h at room temperature followed by incubation overnight at 4°C with a specific primary antibody to p-p53^ser15^ (Cell Signaling, 9284S), β-actin (Cell Signaling, 84575S), Lamin B1(Cell Signaling, 13435S), γ-H2AX (Cell Signaling, 2577S), HSP90 (Cell Signaling, 4877S), HSP70 (Cell Signaling, 4873S), HSP40 (Cell Signaling, 48715), HSF1(Cell Signaling, 12972S), p-HSF1^ser307^ (Abcam, ab47369, HSF^K298 sumo^ (MyBioSource, MBS9211638), p21(Cell Signaling, 2946S), TCF11/NRF1 (Cell Signaling, 8052S), Rpn6/PSMD11 (Cell Signaling, 14303S), Rpt5/PSMC3 (Cell Signaling, 13923S), at a dilution of 1:1,000 in blocking buffer with 0.1% Tween-20, followed by incubation with Donkey anti-Rabbit (Li-Cor Biosciences, 926-32213) or anti-mouse secondary antibody (Li-Cor Biosciences,926-32212) at a dilution of 1:5,000 in blocking buffer. After three washings with PBS, immunoblots were visualized using the Odyssey infrared imaging system (Li-Cor Biosciences).

### Statistical analysis

Statistics were performed using GraphPad Prism 8.0 software (GraphPad, San Diego, CA). Two-group comparisons were analyzed by unpaired Student’s t-test and multiple-group comparisons were performed using one-way analysis of variance (ANOVA) followed by Tukey post hoc analysis. Statistical significance was achieved when P < 0.05 at 95% confidence interval.

## Results

### Cellular proteostasis is impaired in IPF fibroblast

Previous reports have shown that the lung fibroblast in IPF becomes senescent [8-10], thereby providing a model for further mechanistic investigations. To verify these reports, purified preparations of primary lung fibroblasts were isolated from the lungs of normal volunteer (NV) and IPF patients during clinically indicated procedures and examined for the expression of senescence markers. Consistent with published studies, we detected a marked increase in senescence markers in fibroblast cells from the lungs of IPF. This included the cyclin-dependent kinase inhibitors p21, the cell cycle arrest protein p53, the DNA damage marker y-H2AX, and SA-β-gal staining (Fig. 1A-C). Furthermore, transcript levels for multiple senescence-associated secretory proteins, such as *Il6, Mcp1, Mmp2* and *Il1*β and markers of myofibroblast activation (α-Sma, Fn1 and Col1a1) were also increased (Fig. 1D and E).

**Figure 1:**
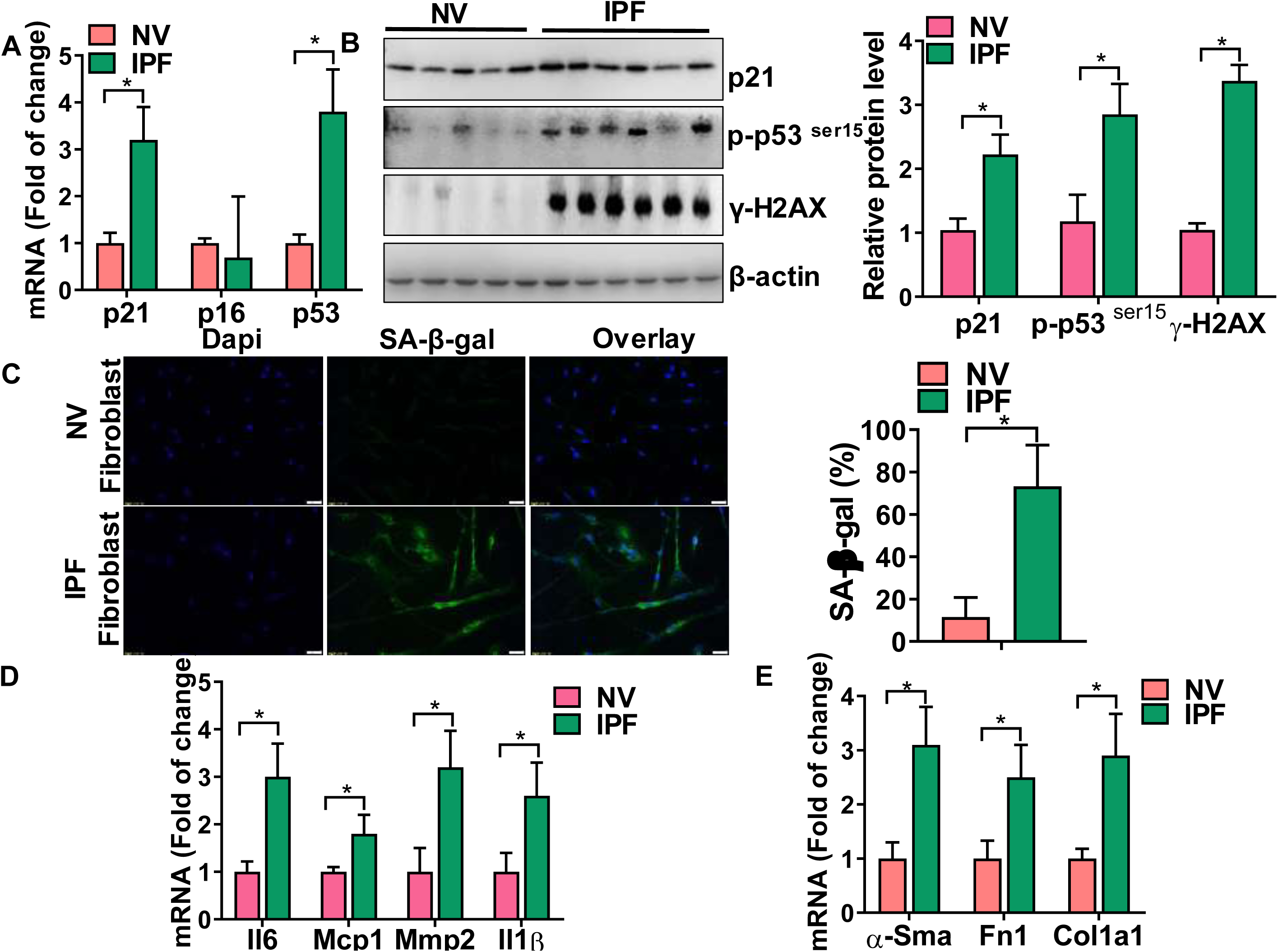
Cellular senescence markers are increased in IPF lung fibroblast. A) Transcript levels for p21, p16, p53 in IPF lung fibroblast and controls. B) Western blot (WB) for p-p53^ser15^, p21 and γ-H2AX in human lung fibroblast cells (with β-actin loading control). Densitometry is shown on the right. C) β-Gal activity (green fluorescence cytoplasmic staining) in IPF and control fibroblast cells. Number of β-Gal positive cells per 100 cells counted (right). D, E) Transcript levels for Il6. Mcp1, Mmp2, Il1β, α-Sma, Fn1 and Col1a1 in human lung fibroblast from IPF and controls. WB images are representative of two different blots. Statistical significance was assessed by Student t-test * p< 0.05 versus control group, n=5-6.

We next evaluated whether proteosasis network was altered in these cells. First, we focused on HSP expression since regulation of these chaperones is believed to be the major contributor of cytoplasmic proteostasis and reduced expression of HSP72 has been found in IPF fibroblast [11-13]. Consistent with the downregulation of this pathway, we found that gene and protein levels of *Hsp70, Hsp90, and Hsp40* were significantly decreased in fibroblasts from lungs of IPF subjects (Fig. 2A and B). Because the transcription factor HSF1 drives the expression of molecular chaperones, we next sought to determine whether downregulation of this pathway was associated with a decrease in HSPs. Unexpectedly, as shown in Fig 2C, we found that nuclear levels of HSF1 were increased by nearly 2-fold in fibroblasts from the lungs of IPF. Since, HSF1 functional activity can be repressed by post-transcriptional modification, including phosphorylation at serine 307 and sumoylation at K298, we next thought to investigate whether this reduction in the cellular chaperones expression was due to post-transcriptional inactivation of the HSF1. Remarkably, HSF1 phosphorylation at Ser307 and sumoylation at K298 were increased by over 1.5-fold in fibroblasts from the lungs of IPF (Fig. 2C), suggesting a link between impaired HSF1 transactivation, reduction in proteins chaperones, and cellular senescence in these cells.

**Figure 2:**
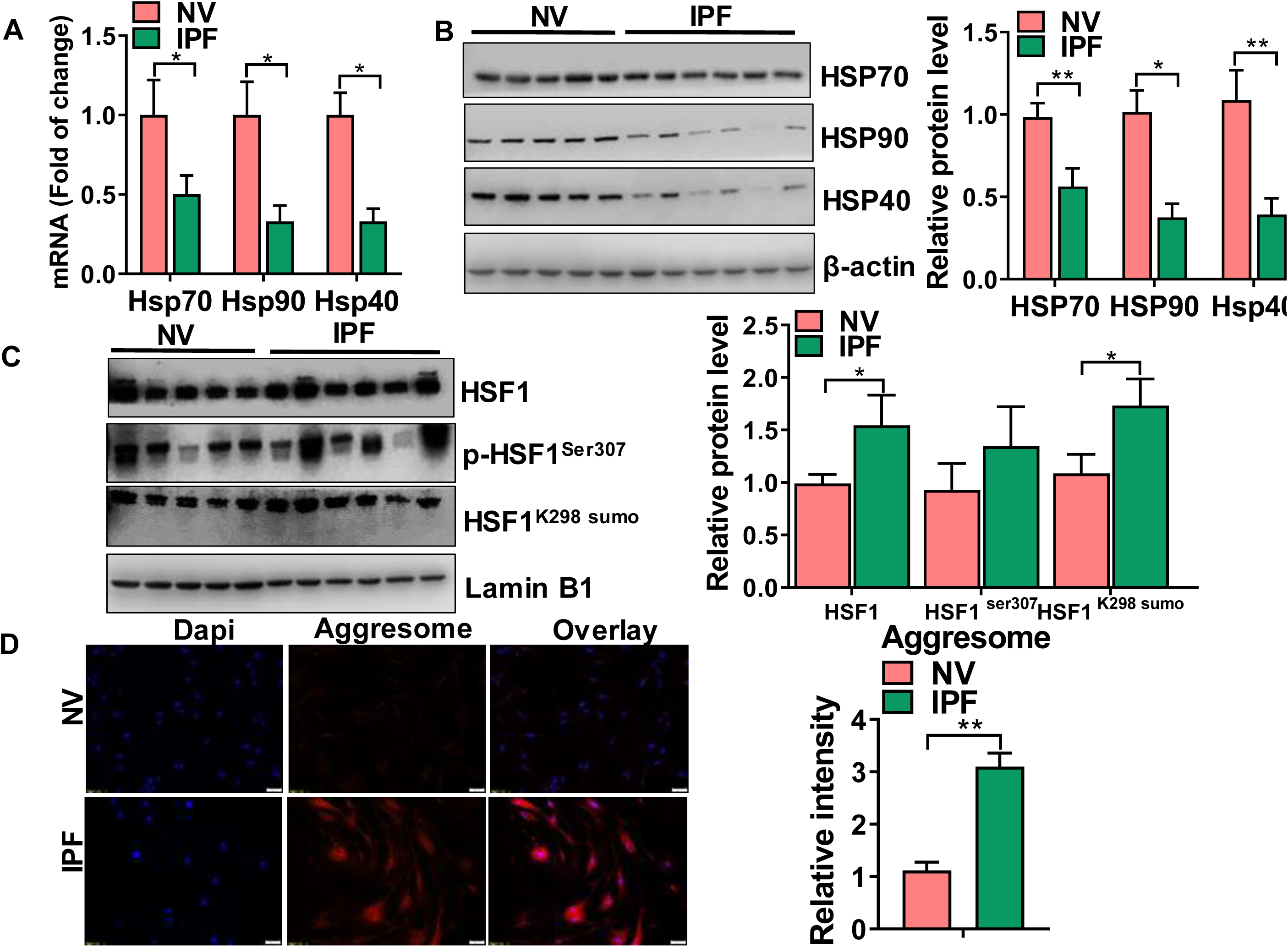
HSF1-mediated induction of chaperones is impaired in IPF lung fibroblast. A) Transcript levels for Hsp70, Hsp90 and Hsp40 in human lung fibroblast from IPF and controls. B, C) WB for HSP90, HSP70, HSP40, HSF1, p-HSF1^ser307^, HSF1^K298 sumo^ in human fibroblast isolated from IPF and controls. D) Quantification of protein aggregation (aggresome) levels by Proteostat staining of IPF lung fibroblast and controls. Western blot images are representative of two different blots and results of densitometry analysis are depicted in bar graphs (n=5-6, per group). Statistical significance was assessed by Student t-test * p<0.05, ** p< 0.01 versus control group.

Next, in order to evaluate whether decline in HSF1 activity and expression of chaperones proteins was associated with a decline in fibroblast proteostasis, we used a recently established molecular rotor to measure intracellular protein aggregation. As expected, we found dramatically increased intracellular protein aggregation in fibroblast isolated from lung of IPF (Fig. 2D).

### Lung fibroblasts isolated from bleomycin injured mice show features of cellular senescence

To evaluate whether senescence markers were upregulated during experimental fibrosis, we measured transcript levels for various senescence markers in isolated murine lung fibroblasts after chronic bleomycin challenge. Strikingly, we detected a marked upregulation in transcript levels for *p21 and p53* in lung fibroblasts of chronic injured mice (Fig. 3A). Similar to IPF fibroblasts, we also found that SA-β-gal staining was increased in lung fibroblasts isolated from the lungs of bleomycin-injured mice relative to age-matched controls (Fig. 3B), and this was associated with a marked upregulating in transcript levels for multiple SASP genes, including as *Il6, Mcp1, Mmp2* and *Il1*β (Fig.3C) and markers of fibroblast activation (Fig. 3D). Consistent with this being a severe pulmonary insult, we found that chronic bleomycin had a dramatically effect on collagen deposition in the lungs of mice at day 60 after injury (Fig. S1). These data demonstrated a substantial induction of senescence marker expression in lung fibroblasts after bleomycin administration.

**Figure 3.**
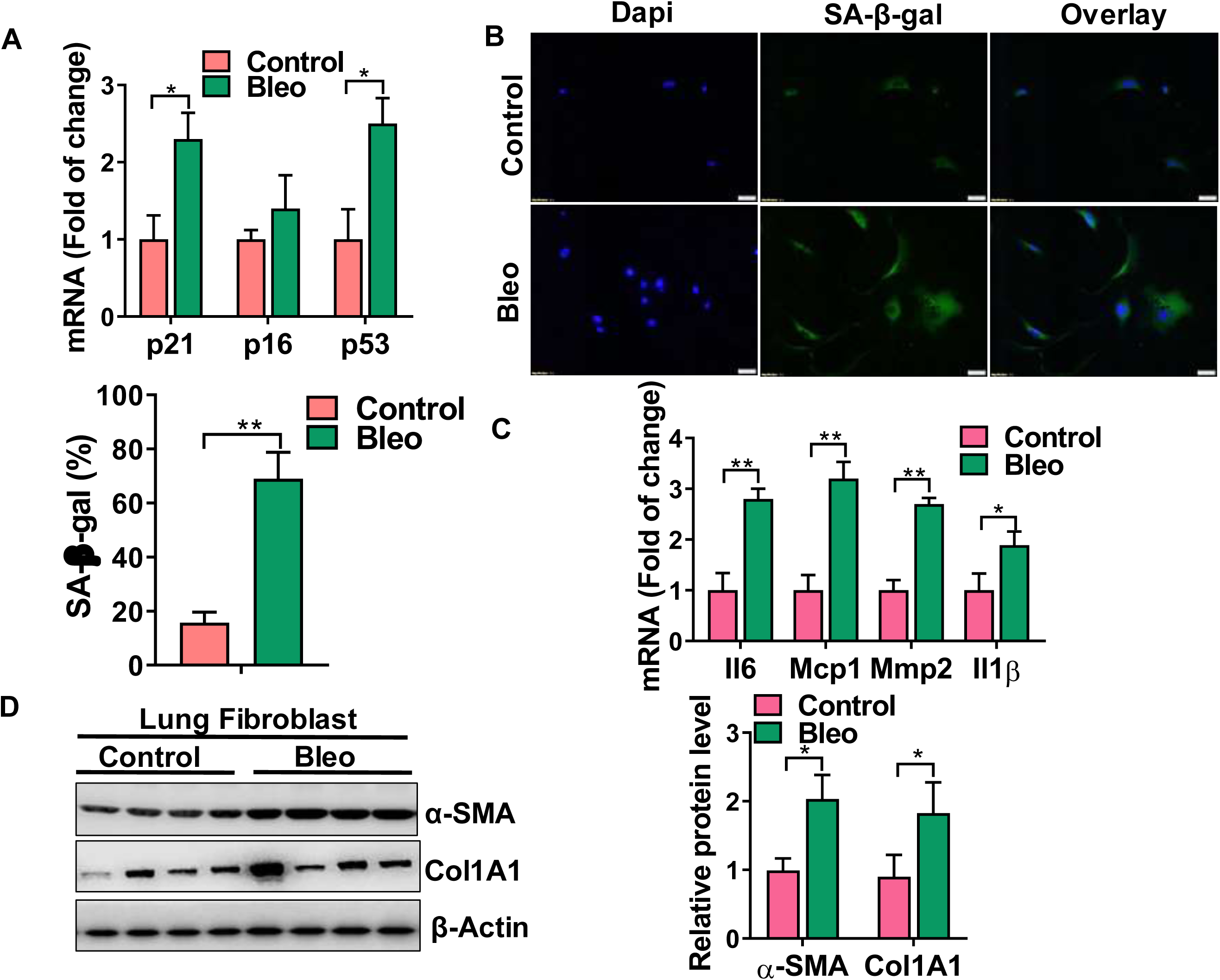
Lung fibroblast isolated from bleomycin-injured mice exhibit features of cellular senescence. A) Transcript levels for p21, p16, p53 in lung fibroblast isolated from mice treated with bleomycin and controls. B) β-Gal activity (green fluorescence cytoplasmic staining) in lung fibroblast from bleomycin and controls. Number of β-Gal positive cells per 100 cells counted (right). C) Transcript levels for Il6, Mcp1, Mmp2, Il1β in lung fibroblast from bleomycin-injured and controls. D) WB for α-SMA and COL1A1 in lung fibroblast from bleomycin and controls. Densitometry is shown on the right. Images are representative of two different blots and results of densitometry analysis are depicted in bar graphs (n= 5-6, per group). Statistical significance was assessed by Student t-test * p<0.05, ** p< 0.01 versus control group.

### HSF-1-mediated chaperone expression is impaired in lung fibroblasts after bleomycin challenge

To assess whether chronic bleomycin challenge alters chaperone levels in the lung fibroblast, we first performed quantitative PCR to evaluate transcript levels for several different chaperones known to be expressed in the mouse lung and which have been linked to the proteostasis network, including *Hsp70, 90 and 40*. As demonstrated in Figure 4A, we found that transcript levels for each of the chaperones were evaluated and readily detectable in the lung fibroblast isolated from bleomycin treated mice. Next, to determine whether decline in HSPs expression was associated with a decline in proteostasis, we evaluated the aggregation of proteins in lung fibroblasts. Reminiscent to the human lung fibroblasts, we detected a marked increase in cytoplasm aggresome in lung fibroblast isolated from bleomycin treated mice (Fig. 4C).

**Figure 4.**
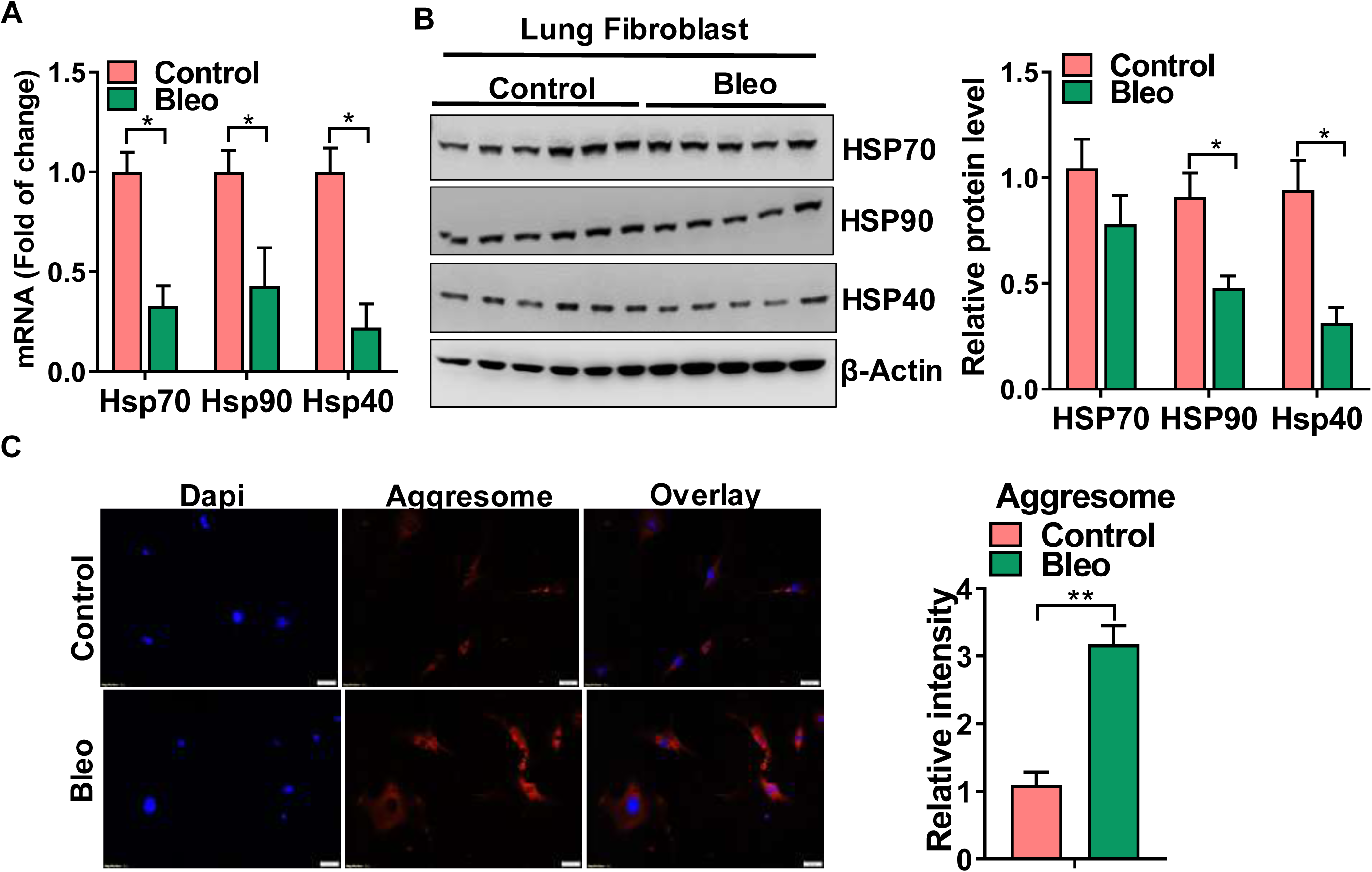
The HSF1 activity is impaired in lung fibroblast isolated from bleomycin-injured mice. A) Transcript levels for Hsp70, Hsp90 and Hsp40 in lung fibroblast isolated from mice treated with bleomycin and controls. B) WB for HSP90, HSP70 in fibroblast cells from bleomycin-injured lungs and controls. Densitometry is shown on the right. C) Quantification of protein aggregation (aggresome) levels by Proteostat staining of lung fibroblast isolated from bleomycin-injured mice and controls. Western blot images are representative of two different blots and results of densitometry analysis are depicted in bar graphs (n=5-6, per group). Statistical significance was assessed by Student t-test * p<0.05, ** p< 0.01 versus control group.

### Characterization of oxidant-induced senescence in cultured lung fibroblasts

To further explore the relationship between cellular senescence and proteostasis, we next developed an *in vitro* model system that would enable more detailed mechanistic investigations. We started by testing various concentrations and exposures to H_2_O_2_, a chemical oxidant previously shown to induce senescence in fibroblasts [38]. Afterwards, we found that cellular senescence could be readily induced in mouse lung fibroblast after 5 consecutive days of H_2_O_2_ (10 µM) exposure (Fig. 5A). This was evident by not only upregulation in numerous cellular senescence markers, including p-p53, p21, y-H2AX and the senescence-associated secretory proteins *Il6, Mcp1, Mmp2* and *Il1*β (Fig. 5B-D), but also by an increase in SA-β-gal staining (Fig. 5E). Considering the similarities identified in these cells when compared to IPF cells, further experiments were conducted in H_2_O_2_-treated lung fibroblasts to identify mechanism of action.

**Figure 5.**
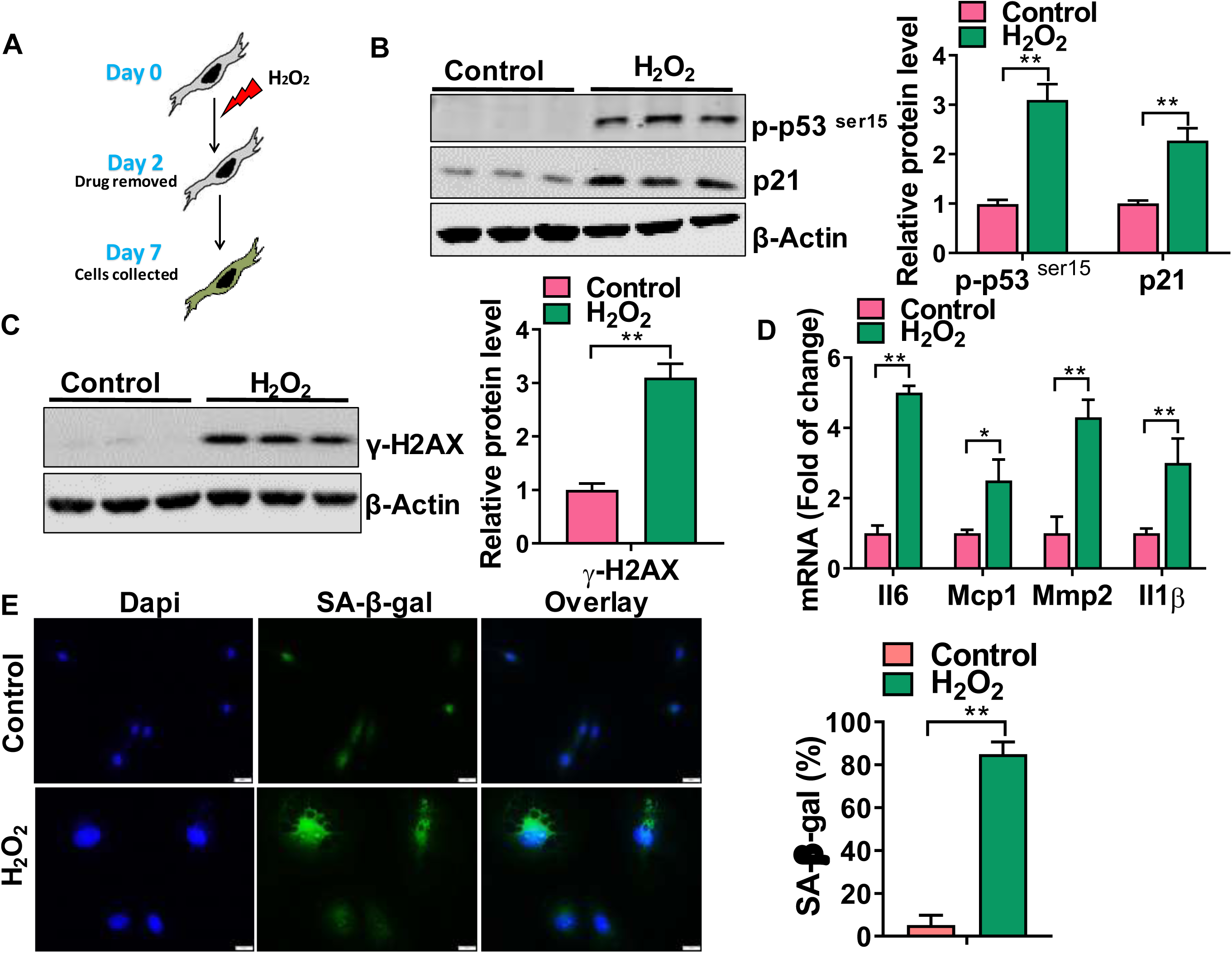
Chronic low dose of H_2_O_2_ induces senescence in mouse Mlg fibroblast cells. A) Schematic showing the experimental design: Mlg cells were treated with H_2_O_2_ (10 μM) for 5 consecutive days. The culture medium was removed and cells were maintained in culture for an additional 2 days. B, C) Western blot (WB) for p-p53^ser15^, p21 and γ-H2AX levels in control and senescent Mlg cells. D) Transcript levels for Il6. Mcp1, Mmp2, Il1β in control and senescent Mlg cells. E) β-Gal activity in control and senescent Mlg cells. Number of β-Gal positive cells per 100 cells counted (right). Densitometry is shown on the righ. Images are representative of two different blots and results of densitometry analysis are depicted in bar graphs (n=6, per group). Statistical significance was assessed by Student t-test * p<0.05, ** p< 0.01 versus control group.

### Mitochondrial respiration is increased in senescent lung fibroblasts

Previous studies have shown that increase in mitochondrial respiration is associated with cellular senescence [39, 40]. With this in mind, we next assessed whether mitochondrial respiration, a primary driver of ROS production, was increased in H_2_O_2_-treated lung fibroblasts. Here, mitochondrial OCR was measured using the Seahorse Bioanalyzer as previously described [21]. As shown in figure S1, we found that basal and ATP-linked respiration were both significantly increased in senescent cells (Fig. S2A). Moreover, these changes were associated with marked upregulation in extracellular acidification rate (ECAR) between control and senescence lung fibroblast (Fig. S2B), indicating that senescence fibroblasts upregulate both mitochondrial respiration and glycolytic activity.

### Cellular proteostasis decline in senescence lung fibroblasts

Having established a strong *in vitro* model of cellular senescence, we next assessed whether proteostasis network was altered in these cells treated with H_2_O_2_. Again, our initial focus was on the HSF1-mediated chaperone expression signaling given its dramatic downregulation in fibroblast cells isolated from lung of IPF and bleomycin injured mice. Similarly, we found that *Hsp70, 90, and 40* gene levels, as well as protein expression were decreased in senescent cells (Fig. 6A and 4B). Moreover, this was associated with elevated nuclear levels of HSF1 and increase in the phosphorylation of HSF1 at Ser307 and sumoylation at K298 (Fig. 6C and 6D), findings indicative of transcriptional inactivation of HSF1. Furthermore, we found that cytoplasmic misfolded and aggregates of proteins were dramatically increased in senescent fibroblast (Fig. 6E), again suggesting a link between impaired cellular proteostasis and cellular senescence in lung fibroblasts. Because misfolded and aggregate proteins also increases when clearance mechanisms are reduced, we assessed whether the 26S proteasome activity and proteins levels of key proteasome subunits were perturbed in senescent cells. Consistent with an increase in protein aggregation, we found that chymotrypsin-like and caspase activities were significantly reduced when comparing levels in senescent versus non-senescent cells (Fig. S3A-B). Next, in order to evaluate whether reduction in 26S proteasome activity was due to impaired proteasome subunit production, we evaluated the nuclear levels of the transcription factor 11 (TCF11), a transcription factor that controls proteasome subunit expression. As shown in Fig. S3C-D, we found that the gene and protein levels were dramatically upregulated in senescent cells. Consistent with this increase in TCF11 expression, we found that Rpn6 and Rpt5 were markedly increased when comparing senescent to control fibroblasts (Fig. S3E), indicating that the decline in the proteasome function is mediated by different mechanisms. Altogether, these findings indicate that cytoplasmic proteostasis is profoundly altered in senescent lung fibroblasts, perhaps due to combined effects on the chaperones expression and clearance of misfolded proteins from cells.

**Figure 6.**
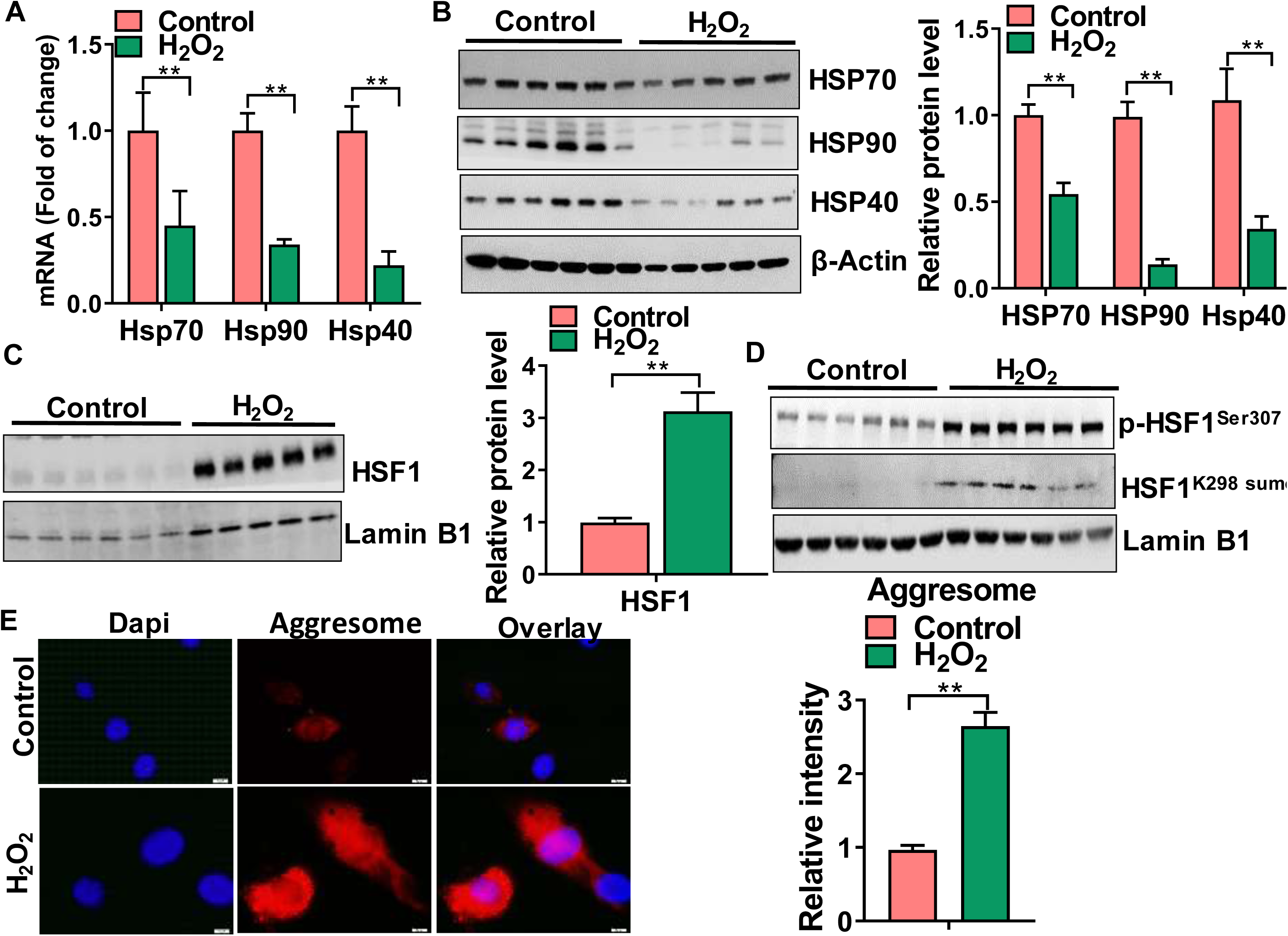
Cellular proteostasis is impaired in senescent Mlg cells. A) Transcript levels for Hsp70, Hsp90 and Hsp40 in control and senescent Mlg cells. B, C, D) WB for HSP90, HSP70, HSP40, HSF1, p-HSF1^ser307^, HSF1^K298 sumo^ in controls and senescent Mlg cells. Densitometry is shown on the right. E) Quantification of protein aggregation (aggresome) levels by Proteostat staining of lung fibroblast controls and senescent. Number of Proteostat positive cells per 100 cells counted (right).Western blot images are representative of two different blots and results of densitometry analysis are depicted in bar graphs (n=6, per group). Statistical significance was assessed by Student t-test * p<0.05, ** p< 0.01 versus control group.

### Small molecule activator of HSF1 increases chaperone expression and reduced cytoplasmic protein aggregation in senescent lung fibroblasts

To test the hypothesis that impaired HSF1 function contributes to proteostasis collapse in our model, we treated H_2_O_2_-exposed fibroblasts with the small-molecule proteostasis regulator A3 (10 µM) in order to activate HSF1 signaling pathway. The A3 molecule belongs to the β-aryl-α,β-unsaturated-carbonyls chemical series and was identified as a new class of small-molecule activators that induced HSF1-dependent chaperones expression [41]. Importantly, treatment with A3 (1 µM-10 µM) did not induce significant toxicity to our cells, as assessed using a standard cytotoxic assay (Data not shown). Consistent with the small molecule activator’s known effects, activation of the HSF1 signaling axis was markedly increased in A3-treated senescent cells, as demonstrated by an increase in HSF1 levels and reduction in the phosphorylation of HSF1 at Ser307 and HSF1^K298 sumo^ expression (Fig. 7A and B). Moreover, this associated with nearly a 2-fold increase in levels of *Hsp70, 90 and 40* gene levels (Fig. 7C), as well as protein expression (Fig. 7D) in A3-treated senescent cells. Also consistent with an increase in HSF1-mediated chaperones expression, we found that cytoplasmic protein aggregation levels were significantly lower in cells treated with A3 (Fig. 7E). More importantly, we found that the small molecule A3 dramatically reduced the expression of cellular senescence markers in H_2_O_2_-exposed cells. This included a reduction in p21 and p-p53 levels, a decrease in SA-β-gal staining, and a decrease in the expression of *Il6, Mcp1, Mmp2* and *Il1*β (Fig. 8A-C).

**Figure 7.**
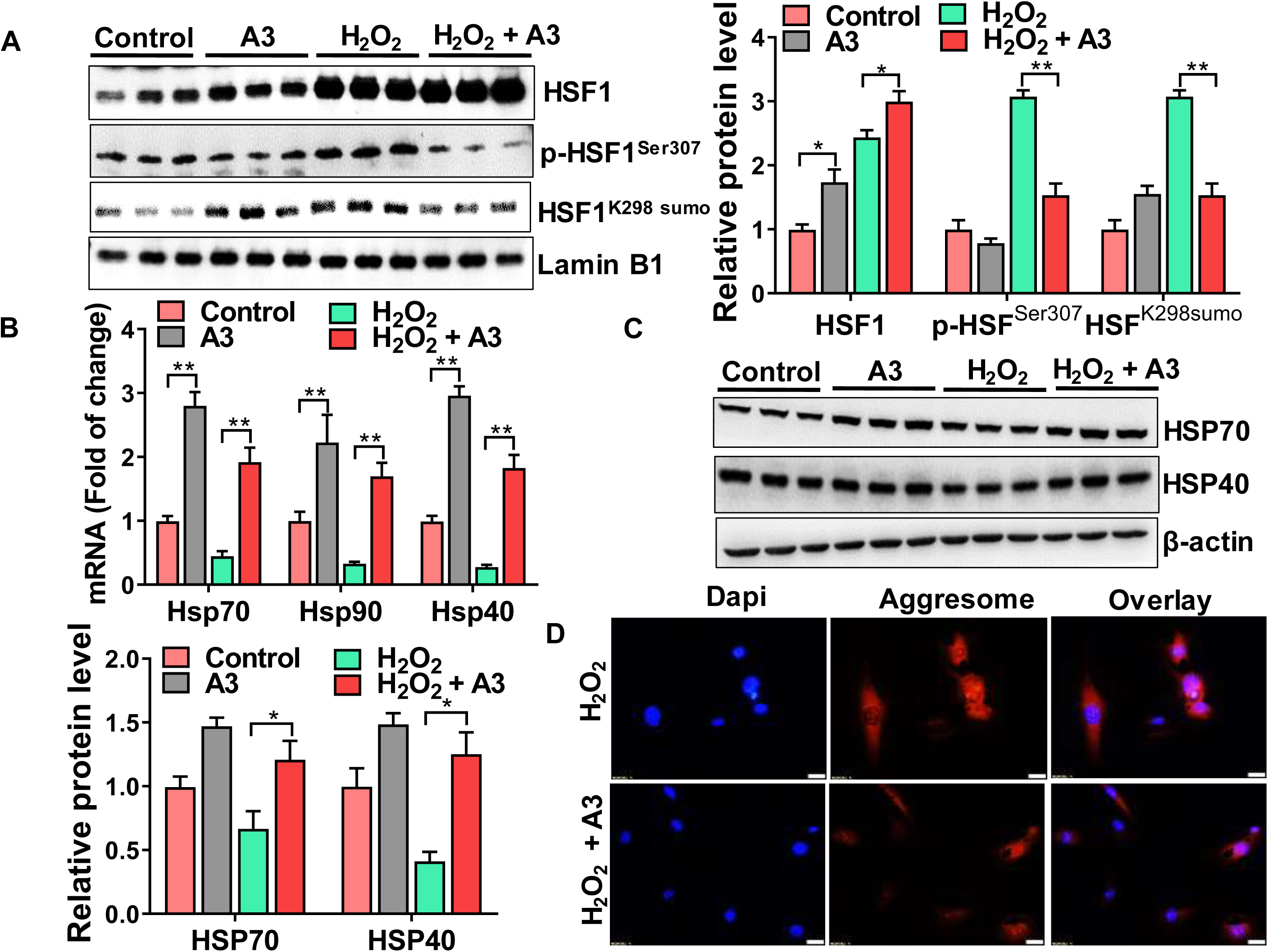
Small molecule activator of HSF1 (A3) restore chaperones expression and reduced cytoplasmic protein aggregation in senescent lung fibroblast cells. A) WB for HSF1, p-HSF1^ser307^, HSF1^K298 sumo^ in control, Mlg+A3, Mlg+H_2_O_2_ and Mlg+H_2_O_2_+A3. B) Transcript levels for Hsp70, Hsp90 and Hsp40 in control, Mlg+A3, Mlg+H_2_O_2_ and Mlg+H_2_O_2_+A3. C) WB for HSP70, HSP40 in control, Mlg+A3, Mlg+H_2_O_2_ and Mlg+H_2_O_2_+A3. D) Quantification of protein aggregation (aggresome) levels by Proteostat staining in control, Mlg+A3, Mlg+H_2_O_2_ and Mlg+H_2_O_2_+A3. Number of Proteostat positive cells per 100 cells counted (right).Western blot images are representative of two different blots and results of densitometry analysis are depicted in bar graphs (n=6, per group). Statistical significance was assessed by ANOVA, * p<0.05, ** p< 0.01.

**Figure 8.**
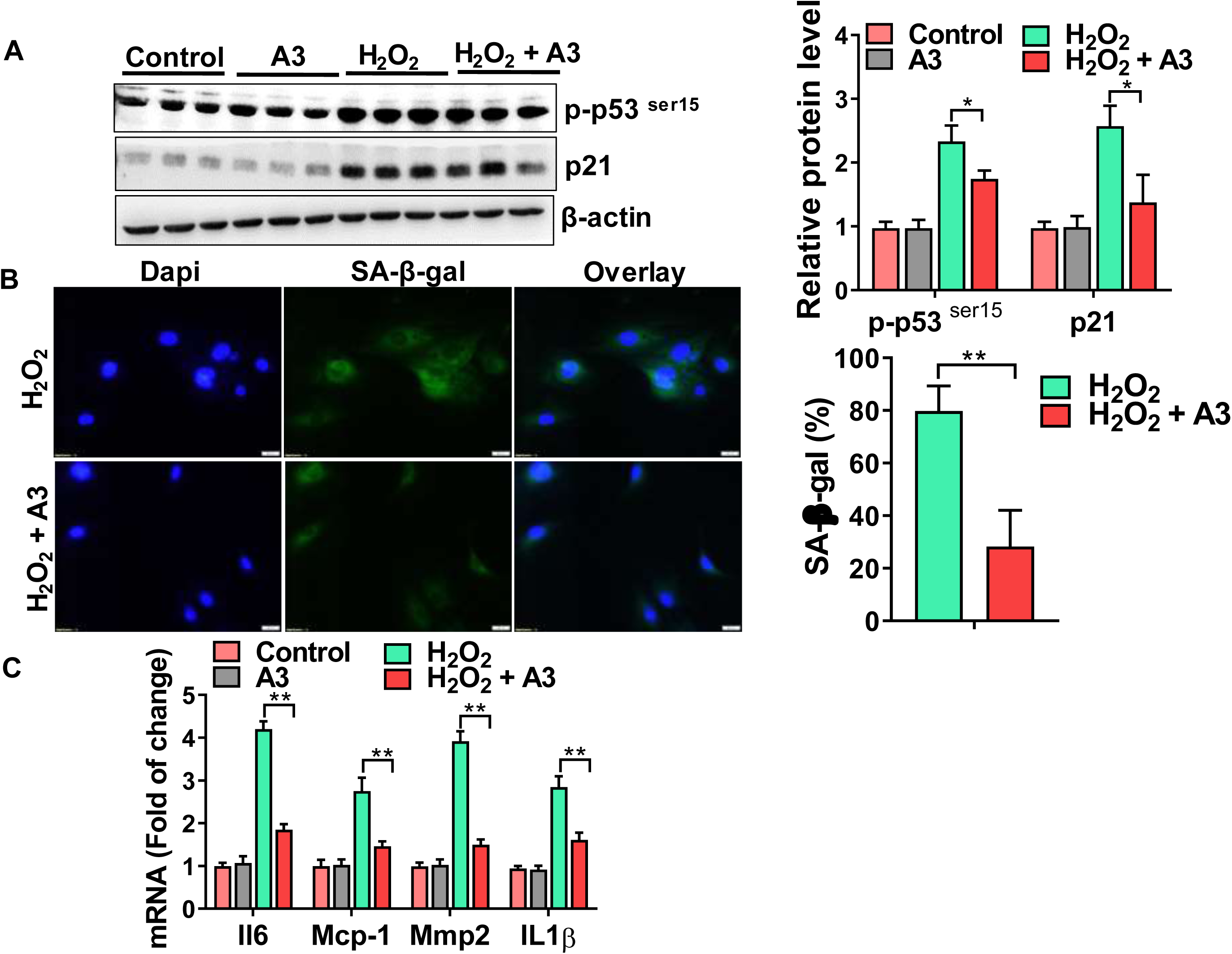
A3 treatment blocks the induction of cellular senescence in H_2_O_2_-exposed mouse Mlg cells. A) Western blot (WB) for p-p53^ser15^, p21 and β-actin in control, Mlg+A3, Mlg+H_2_O_2_ and Mlg+H_2_O_2_+A3. B) Representative images of β-Gal activity (green fluorescent cytoplasmic staining) in control, Mlg+A3, Mlg+H_2_O_2_ and Mlg+H_2_O_2_+A3. Percentage of β-gal stained cells are shown on the right. C) Transcript levels for Il6. Mcp1, Mmp2, Il1β in control, Mlg+A3, Mlg+H_2_O_2_ and Mlg+H_2_O_2_+A3 groups. Western blot images are representative of two different blots and results of densitometry analysis are depicted in bar graphs (n=6, per group). Statistical significance was assessed by ANOVA * p< 0.05, **p<0.01.

### Activation of HSF1-mediated chaperone expression reduces protein aggregation, cellular senescence and pro-fibrotic factors in IPF fibroblasts

Next, to assess whether the small molecule activator of HSF1 can reduce senescence and pro-fibrotic factors, we treated IPF fibroblast with A3. Remarkably, we found that impaired HSF1 transactivation was almost completely abrogated in response to A3 treatment, as demonstrated by the dramatic increase in *Hsp70, 90, and 40* gene levels (Fig. 9A) and reduction in aggregates of proteins (Fig. 9B). Moreover, we found that A3 markedly suppressed the expression of cellular senescence markers (Fig. 10A and 10B). However, activation of HSF1-mediated chaperone expression does not affect the transcript levels of SASP factors (Fig. 10C). Interestingly, A3 markedly suppressed the expression of myofibrobast activation markers in IPF lung fibroblasts (Fig. 10D), suggesting that HSF1 dysfunction and the consequent protein misfolded/aggregation accumulation might be upstream of IPF myofibroblast activation.

**Figure 9.**
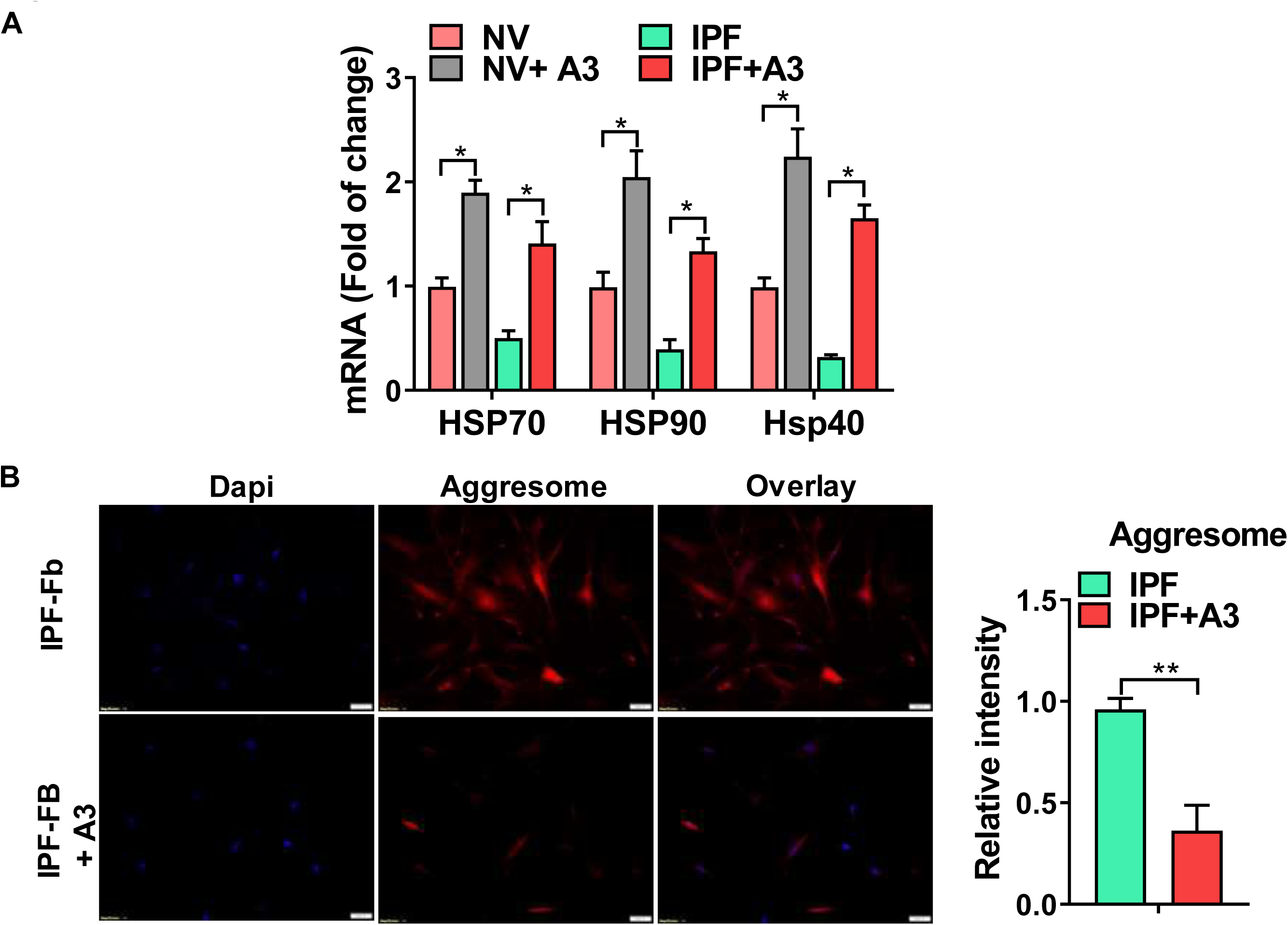
A3 treatment increases HSPs expression and reduced protein aggregation in IPF fibroblast cells. A) Transcript levels for Hsp70, Hsp90 and Hsp40 in normal volunteers (NV), NV+A3, IPF and IPF+A3. B) Quantification of protein aggregation (aggresome) levels by Proteostat staining in NV, NV+A3, IPF and IPF+A3. Number of Proteostat positive cells per 100 cells counted (right). Statistical significance was assessed by ANOVA * p< 0.05, **p<0.01.

**Figure 10.**
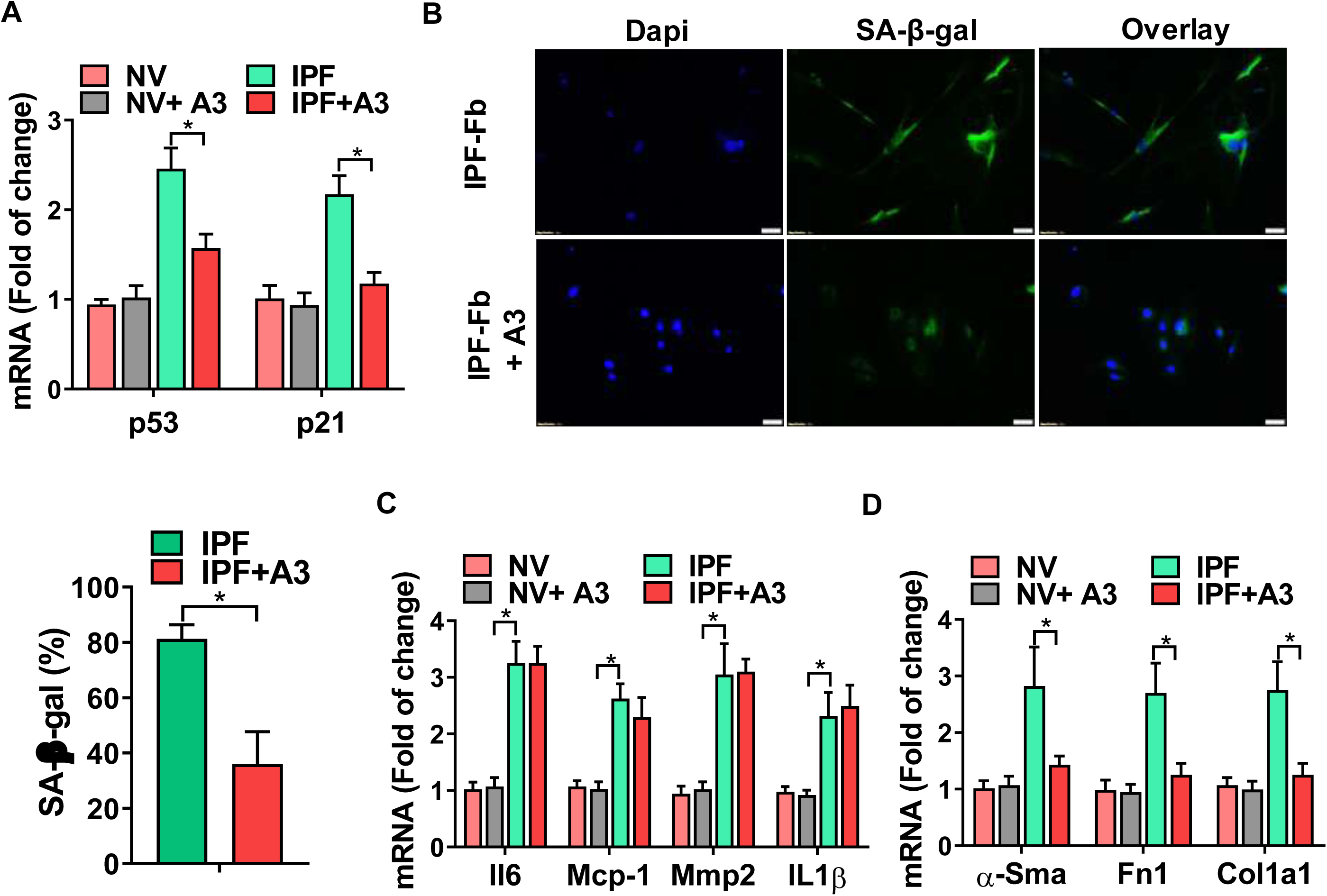
A3 treatment reduces cellular senescence markers and myofibroblast activation in IPF fibroblast. A) Transcript levels for p53, and p21 in normal volunteers (NV), NV+A3, IPF and IPF+A3. B) Representative images of β-Gal activity in normal volunteers (NV), NV+A3, IPF and IPF+A3 groups. Percentage of β-gal stained cells are shown on the right. C, D) Transcript levels for for Il6. Mcp1, Mmp2, Il1β, α-Sma, Fn1, Col1a1. Statistical significance was assessed by ANOVA * p< 0.05.

## Discussion

Cellular senescence and decline in HSP70 expression of lung fibroblasts has emerged as an important contributor to the development of pulmonary fibrosis [11-13]. Fibroblast senescence is thought to cause lung pathology through various mechanisms, including promoting lung inflammation and fibrosis through the secretion of mediators and extracellular matrix proteins [42]. Here, our objective was to elucidate the mechanisms by which HSPs decline and their relationship with cellular senescence in lung fibroblasts. By using different approaches, we show that senescent lung fibroblasts exhibit an increase in HSF1 phosphorylation at Ser307 and sumoylation at K298 and this is associated with a downregulation in protein chaperone expression and the accumulation of protein aggregates. Furthermore, our mechanistic studies support the notion that the decline of this pathway plays a causal role in inducing cellular senescence and fibroblast activation. The data suggest that targeting the HSF1-dependent chaperone expression in lung may represent a novel way to reduce the onset or severity of senescence-associated respiratory diseases.

The principal target of HSF1 are genes encoding molecular chaperones, primarily belonging to HSP families including HSP90, HSP70, HSP40 and small HSPs as well as proteins involved in the regulation of cell metabolism, growth and survival [43]. More than 13 different HSP genes have been identified within the mouse genome and even more are believed to exist in humans [44]. HSPs are important in restoring misfolded conformation and prevent excessive protein denaturation and aggregation. HSF1 deficiency has been reported to induce cellular senescence, affecting numerous signaling pathways including redox homeostasis, antioxidant defense and p53 signaling [31, 32]. In this study, we used a targeted approach to examine HSPs levels in the lung fibroblast, measuring only changes in three key HSPs; HSP70, HSP90, and HSP40, known to be expressed in mouse and human respiratory tissues. Here, we uncovered the downregulation of several HSPs in the lung fibroblast of IPF. Interestingly, we also detected a downregulation in different HSPs transcripts and protein levels in the lungs of bleomycin-injured mice and in our *in vitro* model, including a greater than 2-fold decrease in levels for Hsp70, Hsp90, and Hsp40. Importantly, these findings support the current paradigm in the aging field that proteostasis collapse contribute to cellular senescence [22-24].

Our data agree with recent finding showing that HSPs expression is greatly compromised in cellular senescence [45]. This connection is probably best exemplified in the aging cardiovascular system in which senescence human coronary artery endothelial cells have diminished heat shock response and impaired proteostasis [46]. Proteostasis collapse has also been described in the lungs of patients with various other respiratory conditions including COPD and IPF. For example, the human bronchial epithelial cell line Beas2 and murine lungs exposed to cigarette smoke extract showed a significantly impaired autophagy pathway, cellular senescence and accumulation of aggresome bodies. Moreover, this work also found a significant increase in levels of aggresome bodies in the lungs of smokers and COPD subjects in comparison to nonsmoker controls. The presence and levels of aggresome bodies statistically correlated with severity of emphysema and alveolar senescence [47]. Relevant to our work, researchers have also shown that HSP70 levels is reduced throughout the lung of patients with IPF. Interestingly, Hsp70-knockout mice exposed to a one-time dose of bleomycin into the oropharynx which demonstrated accelerated fibrosis compared to wild-type controls mice [12]. In addition, Sibinska et al., recently demonstrated that HSP90 has an important role in TGF-β1 signaling and that HSP90 inhibition by 17-N-allylamino-17-demethoxygeldamycin (17-AAG) attenuated fibroblast activation and reduce pulmonary fibrosis in mice [48]. However, to our surprise, we found that levels of HSP90 was markedly downregulated in IPF fibroblasts as well as in the lung fibroblasts isolated from bleomycin-injured mice. Taken together, these findings indicate that HSPs genes are important for regulating cellular proteostasis in the lung, and suggest that altered chaperones expression due to HSF1 dysfunction might contribute to lung disrepair and fibrotic remodeling.

Recent studies have shown that protein aggregation accelerates the functional decline of different tissues [49]. In support of this idea, we detected a marked increase in protein aggregation in lung fibroblasts of IPF and chronically injured mice and this finding was associated with increases in cellular senescence and myofibroblast activation markers. Although these findings suggest a potential role for blocking protein aggregation in the treatment of age-related lung diseases, we recognize that these concepts have yet to demonstrate clear benefits in the treatment of any age-related.

We observed that treating lung fibroblasts with the HSF1 activator, A3, reduced protein aggregation and senescence myofibroblast phenotype. We reasoned that, because A3 enhances the expression of multiple molecular chaperones, this treatment might more effectively drive global HSPs expression compared to strategies that focus on any single molecular chaperone. While A3 molecule has known antioxidant properties [41] that may have contributed to its ability to mitigate senescent phenotype, our data indicate that an important feature of the response to A3 was enhanced expression of protein chaperones in the lung fibroblast. In future studies, it will be important to clarify the precise mechanisms by which A3 mediates its effects and to test the ability of HSF1 activators, or other compounds that induce HSPs, to inhibit cellular senescence in related model systems.

Although we postulate that small molecule HSF1 activators could be effective in the treatment of pulmonary fibrosis by increasing HSF1 homotrimerization, resisting stress and antagonizing protein misfolding, we recognize that HSF1 regulation is a complex and multi-step process, and that determining the upstream molecular events leading to its activation has represented a challenge. For example, Qu et al showed that HSF-AB, a mutant form of HSF1 that is unable to associate with HSPs gene promotors, protects neurons from death caused by the accumulation of misfolded proteins [50]. Furthermore, Zou et al demonstrated that HSF1 overexpression significantly attenuated pressure overload-induced cardiac fibrosis by blocking Smad3 activation [51]. While similar strategies were not tested in our study, we anticipate that off-target effects of small molecule activators could be minimized by similar approaches. Contrasting with these benefits, Sasaki et. al., reported that TGF-β and IL-1 induced trimer formation of HSF1, which in turn bound to HSE of HSP47, resulted in the enhancement of HSP47 expression. Furthermore, because HSP47 acts as a universal molecular chaperone for collagen, they proposed that inhibition of HSP47 could be a good candidate for controlling tissue fibrosis [52]. Conversely, several studies have showed an unexpected pro-oncogenic role of HSF1 [53, 54]. HSF1 remains latent in primary cells without stress, it becomes constitutively activated within malignant cells, rendering them addicted to HSF1 for their growth and survival. These finding further highlight the HSF1-mediated HSPs expression as an integral component of the oncogenic network.

In this study, we propose that chaperone proteins were reduced in senescent lung fibroblasts due to an increase in p-HSF1^ser307^ and HSF1^K298 sumo^ levels. However, we recognize that the underlying molecular mechanism remains incomplete. For example, HSF1 phosphorylation at Ser230, Ser419, Thr142, Ser320 and Ser326 are important for its transcriptional activation. By contrast, the inhibitory events include Ser303, Ser307, Ser363 and Ser121 phosphorylation. Several kinases have been implicated in phosphorylation of HSF1 at serine 307, including the MAPK kinase MEK and the p38 MAPK. Notably, among the p38 MAPK family members, p38d MAPK is particularly efficient catalyst of phosphorylation, whereas p38c MAPK is the most specific [33, 34]. Nonetheless, it remains unclear how these events regulate HSF1 activity in IPF fibroblasts. Importantly, HSF1 acetylation can also decrease its transcriptional activity in human cells. Consistent with this, Purwana, et al. recently reported that glucolipotoxicity promoted HSF1 acetylation, inhibits HSF1 DNA binding activity and decreased the expression of its target genes in human beta cells [55]. Interestingly, and similar to our results, these investigators reported that HSF1 total expression was not affected in their model.

In conclusion, this study demonstrates that decline of HSF1 activation plays a central role in causing cellular senescence of the lung fibroblast and supports the concept that this pathway might represent a reasonable target in the treatment of age-related lung diseases.

## Acknowledged

We thank Dr. David Nunley from the Ohio State University, Columbus, OH for collecting and providing subject specimens. We like to acknowledge the support of NIH. Grant No: R01HL136833 (FR).

