## Supporting figures for "Impaired HSF1 transactivation drives proteostasis collapse and senescent phenotype of IPF lung fibroblast"

| Mouse | | |
|---|---|---|
| **Il6** | (F): TCCTTAGCCACTCCTTCTGT | (R): AGCCAGAGTCCTTCAGAGA |
| **I1β** | (F): TGCCACCTTTTGACAGTGATG | (R): TGATGTGCTGCTGCAGATT |
| **Mcp1** | (F): AACTACAGCTTCTTTGGGACA | (R): CATCCACGTGTTGGCTCA |
| **Mmp2** | (F): CAACGATGGAGGCACGAGTG | (R): GCCGGGGAACTTGATCATGG |
| **Hprt1** | (F): AGCAGGTCAGCAAAGAACT | (R): CCTCATGGACTGATTATGGACA |
| **p21/Cdkn1a** | (F): CGAGAACGGTGGAACTTTGAC | (R): CAGGGCTCAGGTAGACCTTG |
| **p16/Cdkn2a** | (F): CTCTGCTCTTGGGATTGGC | (R): GTGCGATATTTGCGTTCCG |
| **Hsp70/Hspa1a/b** | (F): ATGGACAAGGCGCAGATCC | (R): CTCCGACTTGTCCCCCAT |
| **Hsp40/Dnajb1** | (F): CCCCATGCCATGTTTGCT | (R): GCGCTGCCCAAAAAAGG |
| **Hsp90/Hsp90aa1** | (F): GGGCCCGCCTCTATATAAGG | (R): GACCTCCTCCTCCTCCATTG |
| **p53/Trp53** | (F): TTATGAGCCACCCGAGGC | (R): GTACGGCGGTCTCTCCCAG |

| Human | | |
|---|---|---|
| **Il6** | (F): CGGCTACATCTTTGGAATCTTC | (R): GCCCAGCTATGAACTCCTTC |
| **Il1β** | (F): CTCGCCAGTGAAATGATGGCT | (R): GTCGGAGATTCGTAGCTGGAT |
| **Mcp1** | (F): GCCTCTGCACTGAGATCTTC | (R): AGCAGCCACCTTCATTCC |
| **Mmp2** | (F): GGAATGCCATCCCCGATAAC | (R): CAGCCTAGCCAGCCAGTCGGATTT |
| **Gapdh** | (F): GGAAGGGCTCATGACCACAG | (R): ACAGTCTTCTGGGTGGCAGTG |
| **p21/Cdkn1a** | (F): GAGACTAAGGCAGAAGATGTAGAG | (R):GCAGACCAGCATGACAGAT |
| **p16/Cdkn2a** | (F): TGAGCTTTGGTTCTGCCATT | (R): AGCTGTCGACTTCATGACAAG |
| **Hsp70/Hspa1a/b** | (F): GCCTTTCCAAGATTGCTGTT | (R): TCAACATTGCAAACACAGGA |
| **Hsp40/Dnajb1** | (F): CTTTCCCCAAGAAGGAGAC | (R): ATACGACGGGTATCGTCCTG |
| **Hsp90/Hsp90aa1** | (F): AGAGCCGAGCCGACAGAG | (R): CACCTTGCCGTGTTGGAA |
| **p53/Trp53** | (F): GACGGTGACACGCTTCCCTGGATT | (R): GGGAACAAGAAGTGGAGAATGTCA |
| **Acta2** | (F): GTGAAGAAGAGGACAGCACTG | (R): CCCATTCCCACCATCACC |
| **Fn1** | (F): TGTCAGTCAAAGCAAGCCCG | (R): TTAGGACGCTCATAAGTG TCACCC |
| **Col1a1** | (F): AAGGGACACAGAGGTTTCAGTGG | (R): CAGCACCAGTAGCACCATCATTTC |
| **18S rRNA** | (F): GGCCCTGTAATTGGAATGAGTC | (R): CCAAGATCCAACTACGAGCTT |

**Table Supplementary 1:** Primers used for qRT-PCR.

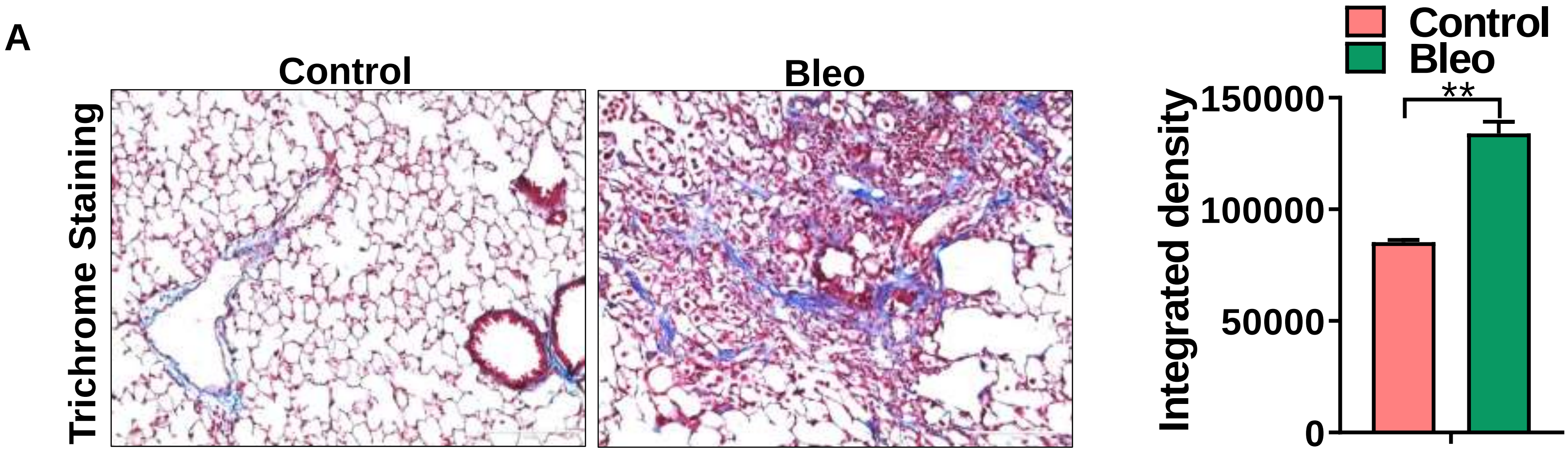


**Figure Supplementary 1:** Low-power view of trichrome staining lung tissue (blue, collagen, 20X) in lung tissues from mice controls and challenged with bleomycin. Densitometry analysis are depicted in bar graphs. Scale bars: 100 µm. Statistical significant was assessed by Student t-test *** $p < 0.001$ versus control group.

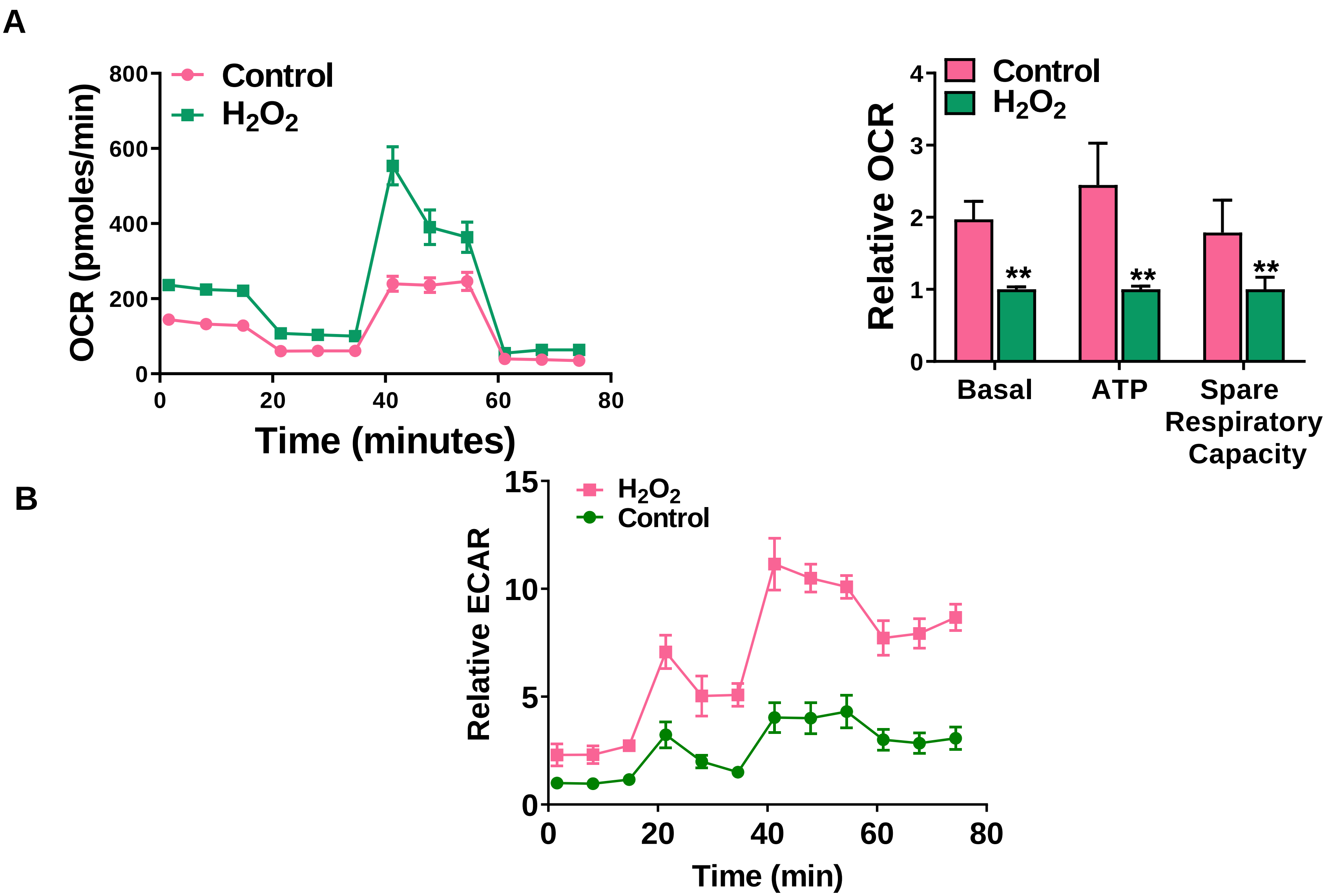


**Figure Supplementary 2:** Oxidative phosphorylation is increased in senescent Mlg cells**.** A) Oxygen consumption rate in control and senescent cells. B) ECAR values in control and senescent Mlg cells. Seahorse data are representative of three separate experiments with control and senescent cells. Statistical significance was assessed by Student t-test $^{**}p<0.01$ versus control group.

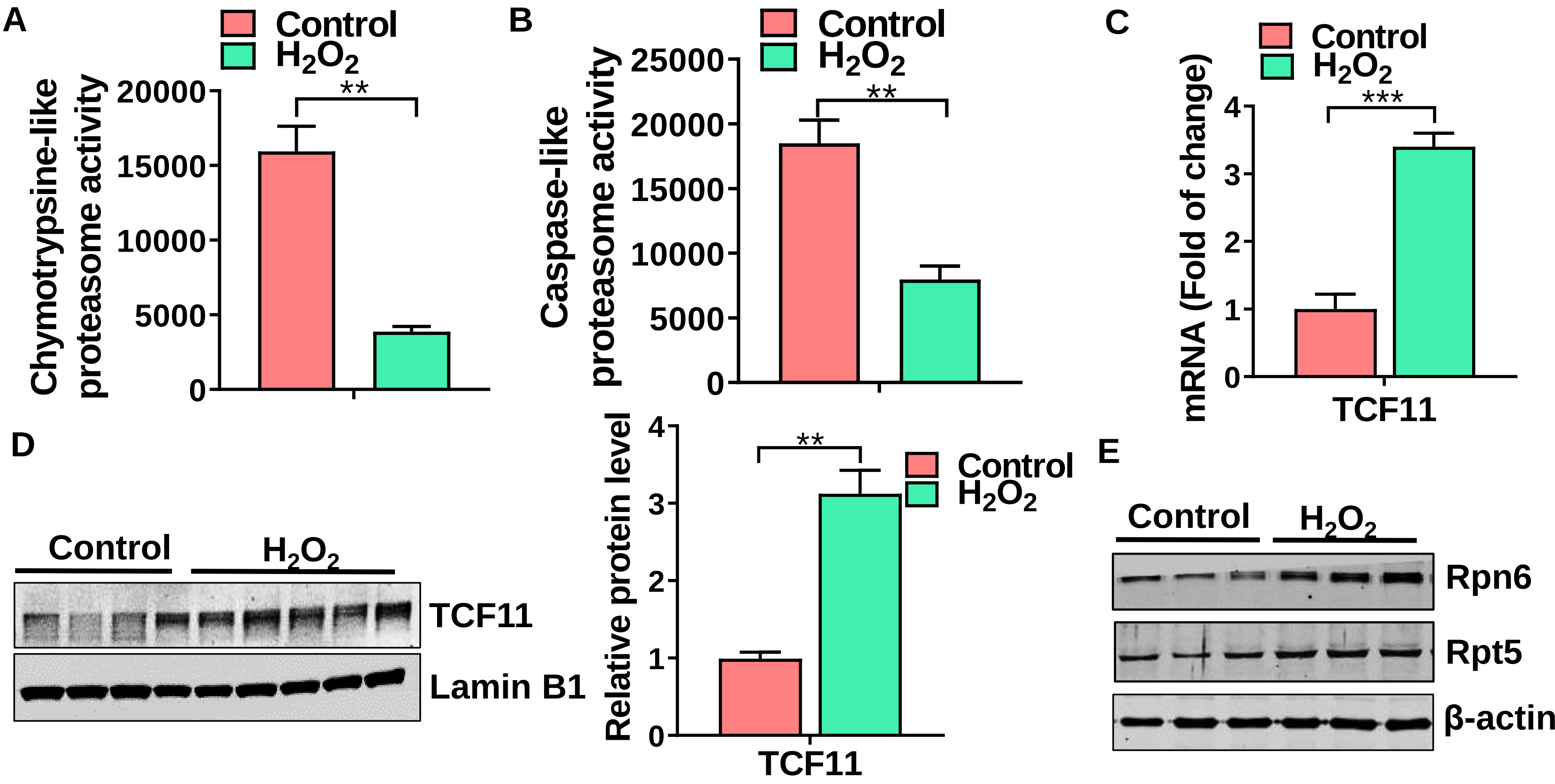


**Figure Supplementary 3:** 26S proteasome activity is impaired in senescent Mlg cells**.** A,B) Chymotrypsin-like and caspase-like activities in control and senescent cells. C) Transcript levels of TCF11 in control and senescent Mlg cells. D) Western blot for TCF11, Rpn6 and Rpt5 in control and senescence Mlg fibroblast. Results are representative of three separate experiments with control and senescent cells. Statistical significance was assessed by Student t-test $^{**}p<0.01$, $^{***}p<0.001$ versus control group.
